# Geometry-driven asymmetric cell divisions pattern cell cycles and zygotic genome activation in the zebrafish embryo

**DOI:** 10.1101/2025.02.03.636134

**Authors:** Nikhil Mishra, Yuting I. Li, Edouard Hannezo, Carl-Philipp Heisenberg

## Abstract

Early embryo geometry is one of the most invariant species-specific traits, yet its role in ensuring developmental reproducibility and robustness remains underexplored. Here, we show that in zebrafish, the geometry of the fertilized egg - specifically its curvature and volume - serves as a critical initial condition triggering a cascade of events that exert a lasting influence on development. Specifically, it guides asymmetric cell divisions in the blastoderm, generating radial gradients of cell volume and nucleocytoplasmic ratio. These gradients generate mitotic phase waves, with individual cell cycle periods determined largely cell-autonomously by the nucleocytoplasmic ratio. Modeling and perturbation experiments demonstrate that reducing cell autonomy reshapes these waves, emphasizing the instructive role of geometry-derived volume patterns in setting the intrinsic period of the cell cycle oscillator. Remarkably, in addition to organizing cell cycles, early embryo geometry also spatially patterns zygotic genome activation (ZGA) at the midblastula transition, a key step in establishing embryonic autonomy. Disrupting geometry alters the ZGA pattern and causes ectopic germ layer specification, underscoring its developmental significance. Together, our findings reveal a novel symmetry-breaking function of early embryo geometry in coordinating cell cycle and transcriptional patterning, establishing a blueprint for robust embryogenesis.

## Main

### Cell cycles in the zebrafish embryo are synchronized in a cell-autonomous manner

Developmental reproducibility and robustness are critical for the survival of a species. Understanding the foundations of this robustness is, therefore, a question of fundamental importance and has become a central focus of research. While most studies have concentrated on the genetic and molecular mechanisms underlying development, the potential influence of geometry on developmental robustness has remained largely overlooked.

Recent work using minimal active matter models has revealed that geometry and curvature play a pivotal role in the mechanical response of epithelial sheets^1–5^. These insights can, for example, account for the anisotropic distribution of Myosin II observed in *Drosophila embryos*^6^. Such findings highlight geometry as a key organizer of mechanical force generation during morphogenesis. Yet, whether - and how - geometry governs more complex, large-scale developmental processes, such as embryo patterning and coordinated cell divisions, remains an open question.

Metazoan development begins with a single cell, the zygote, which undergoes several rounds of rapid divisions (cleavages) to generate a large population of cells capable of adopting multiple fates and exhibiting diverse morphogenetic behaviors. In many species, these early divisions are initially highly synchronous. This cell cycle synchrony is best understood in *Drosophila*, where it is driven by the propagation of a chemical wave. In cleavage-stage *Drosophila* embryos, which are syncytial, nuclear cycles synchronize through "sweep waves" - a combination of coupling-dependent trigger waves and coupling-independent phase waves of Cyclin-dependent kinase 1 (Cdk1) activity propagating through the cytoplasm^7^. Similar mechanisms have been observed in *Xenopus* egg extracts, another syncytial system, where nuclear divisions are coordinated by trigger waves of Cdk1 activity^8^. By contrast, in intact *Xenopus* embryos, where cells are fully separated, early cleavage divisions synchronize independently of cell-cell coupling^9^. Notably, across systems, the propagation of cell cycle waves slows progressively with successive cleavage rounds, leading to ‘metasynchronization’, where cell cycles remain spatially patterned before eventually becoming fully desynchronized^7,9,10^.

Initial cleavages in the zebrafish embryo are meroblastic; i.e., the cytoplasm but not the yolk is partitioned through cytokinesis^10^. Cell cycles at this stage occur highly synchronously, to begin with, and gradually desynchronize (metasynchrony) before completely desynchronizing at division round 10^10^. Importantly, while the cleavage-stage cells cycle between only the S and M phases, complete desynchronization around division round 10 is thought to occur mainly due to a heterogeneous introduction of gap phases to the cell cycles^11^. To determine the spatiotemporal pattern of cell divisions in cleavage-stage zebrafish embryos, we recorded time-lapse images of Tg*(actb2:rfp-pcna)* embryos, allowing us to reliably score the onset of mitosis marked by the disappearance of the nuclear RFP-PCNA localization^12^. Consistent with previous reports, we observed three distinct phases with different extents of mitotic synchrony^10,13^. In the first three cell cleavage division rounds, cell cycles exhibited very high synchrony (**synchronous phase**) (Fig. 1B, S1A-B). Thereafter, cell divisions from cleavage rounds four to nine showed slight but clearly detectable delays (**metasynchronous phase**) (Fig. 1B, S1A-B). These delays increased throughout the metasynchronous phase, with cells in division round four dividing with a mean division timing variance of 0.10 min^2^ and in division round 9 with 3.43 min^2^ (Fig. 1B). Notably, as previously reported, these metasynchronous divisions occurred in a spatiotemporally patterned manner, forming a radial ‘mitotic wave’ that originates near the animal pole (AP), where cells divide first, and propagates toward the margin, where they divide last (Fig. 1C, S1C-D)^10,14,15^. Furthermore, consistent with the increasing variance in division timings during the metasynchronous phase, the speed at which this mitotic wave travels gradually decreased over consecutive division rounds (Fig. 1D). After round nine, cells divided largely non-synchronously, marking the onset of the **asynchronous phase**.

**Fig. 1:**
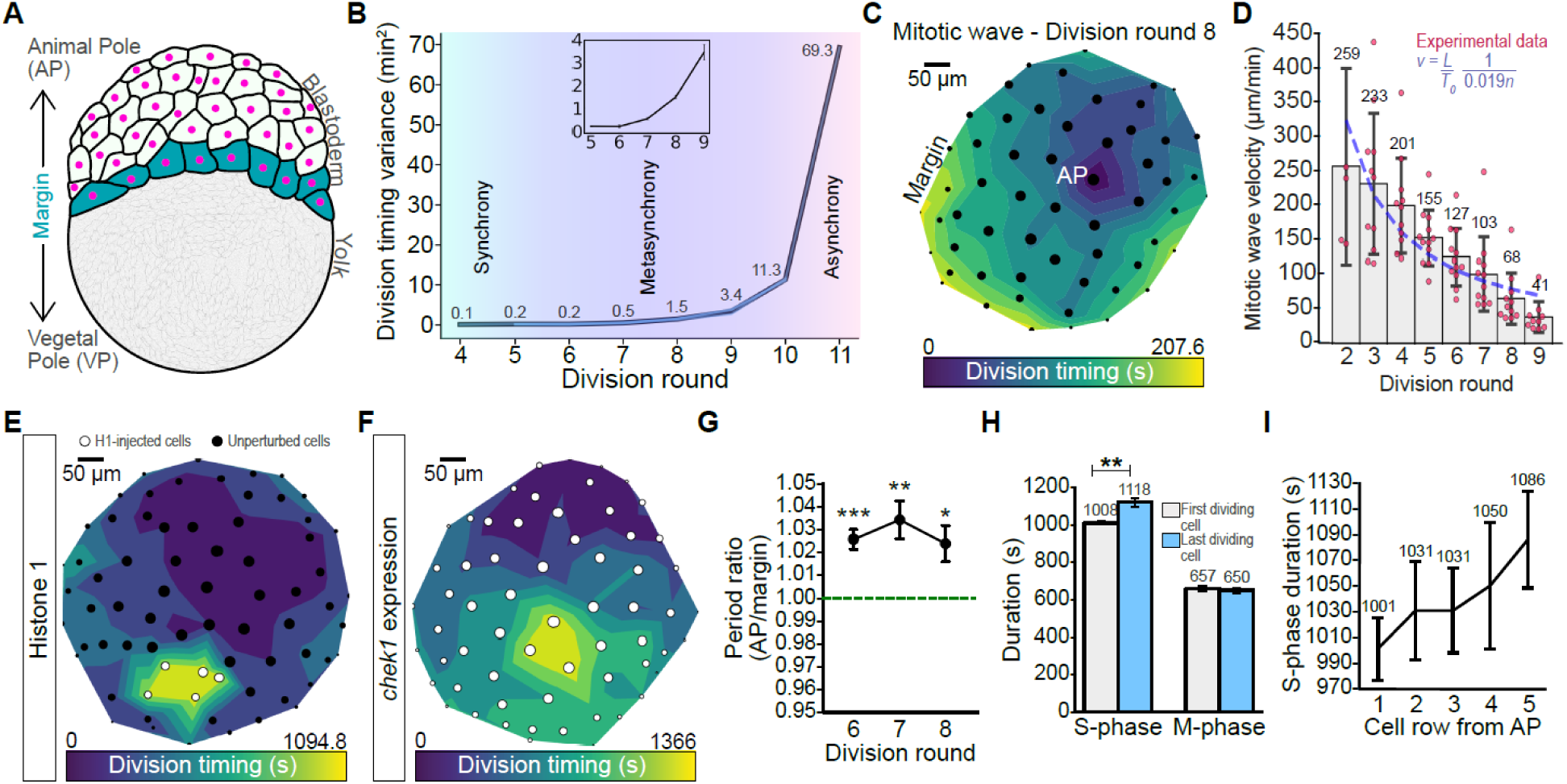
Mitotic synchrony gradually decreases as the S-phase lengthens along the animal-margin axis. (A) Schematic representation of a zebrafish embryo at late cleavage stage. The blastoderm is positioned at the animal pole (AP) atop the yolk cell at the vegetal pole (VP). The last row of blastoderm cells (cyan) constitutes the margin. (B) Line plot of cell division timing variance in Tg(*actb2:rfp-pcna*) embryos for cleavage division rounds 4-11. Blue lines, individual embryos; black line, mean across 4 embryos. Inset shows variance for division rounds 5-9 (C) Contour plot of the radial mitotic wave across all surface cells at the 8th cleavage division round in a representative wild-type embryo. Each dot represents the position of a nucleus along the XY plane, and the size of the dot indicates its position along the Z-axis (AP-margin axis). Colors represent division timing for each nucleus. (D) Bar plot of mitotic wave velocities (mean ± SD), calculated by generating distance-time plots, for cleavage division rounds 2-9, with the red dots showing wave velocities for individual embryos. n=5-11 embryos. Blue dotted line: Curve described by the equation *v*=(*L*/*T0*).(1/0.019*n*), where *L* is the total AP-margin distance, *T0* is the cell cycle period at the AP, and *n* is the division round to represent the effect of a 1.9% period gradient along the AP-margin axis on mitotic wave speeds. (E, F) Representative contour plots of mitotic waves at the 8th round of division in embryos injected with either 0.3pg Histone 1 protein into a single blastomere at the 32-cell stage (E) or 12pg *chek1* mRNA at the 1-cell stage (F). (G) Line plot showing the ratio of cell cycle periods at the AP and margin for division rounds 6-8 (mean ± SEM). n=9 embryos; One sample t-test; *: p<0.05; **: p<0.01; ***: p<0.001. (H) Bar plot of M-phase and S-phase lengths (mean ± SD) at the 8th cleavage division round. n=8 embryos; Student’s t-test with p<0.01]. (I) Line plot of S-phase duration (mean ± SD) in division round 8 as a function of cell position relative to the AP. n≥4 embryos.

To understand how cell cycles are seemingly coordinated into radial mitotic waves during the metasynchronous phase, we asked if their synchronization occurs via cell-cell coupling. To that end, we artificially desynchronized cell cycles at the 32/64-cell stage through mosaic Histone 1 protein injections, previously shown to effectively alter cell cycle progression^16–18^. We found that Histone 1-positive and thus desynchronized cells failed to resynchronize with their neighboring control cells by division round 9, after which cell cycles normally desynchronize in control embryos (Figs. 1E, S2, Movie S2). To address the possibility that these cell cycles could have resynchronized given sufficient time, we assessed cell cycle progression in embryos ectopically expressing the Cdk1 inhibitor, Chek1, through *chek1* mRNA injection in the one-celled embryo, which, by being heterogeneously distributed in the embryo, has previously been shown to interfere with cell cycle synchronization^19^. Chek1-overexpressing embryos exhibited cell cycle desynchronization significantly earlier than control embryos, and failed to resynchronize despite completing all cleavage division rounds (Fig. 1F, S2, Movie S3). Collectively, these observations suggest that cell-cell coupling plays only a minor role, if any, in cell cycle synchronization in the zebrafish embryo.

To determine by which coupling-independent mechanism the radial mitotic wave is formed, we analyzed the evolution of the mitotic wave through consecutive division rounds. We found that the transition from synchrony to metasynchrony was gradual rather than abrupt, suggesting a progressive accumulation of delays rather than distinct switches leading to the observed loss of synchrony (Fig. 1B). Importantly, this gradual loss of synchrony was due to unequal cell cycle periods. For instance, between the metasynchronous division rounds 6-8, cells at the margin cycled with 2-4% longer periods than those at the AP (Fig. 1G). This discrepancy arose exclusively due to a longer S-phase, which increased in cells as a function of their distance from the animal pole (Fig. 1H-I). Although noisy, such period gradients may lead to the observed gradual slowing down of the radial mitotic wave by accumulating increasing delays in the marginal cells over consecutive division rounds. To test this hypothesis, we assumed that the period, *T*, is a linear function of the distance ‘z’ from the AP where 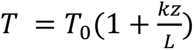, with *T_0_* being the typical period at the AP, *L* the total AP-margin distance, and *k* a dimensionless constant of proportionality. This implies that the speed of the mitotic wave, *v,* in the *n*^th^ division round is expected to be inversely proportional to *n* (Fig. 1D) with 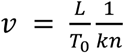. To compare our experimental observations with these theoretical predictions, we first performed linear regression on the division timings to obtain the wave speed for each division round (Fig. S1D). We then fitted the obtained *v* against 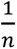 using the least squares fit. Using *k=1.9%,* we obtained a close match between experiments and theory (Fig. 1D), suggesting that a 1.9% slower cell cycle at the margin would, in principle, be sufficient to explain the observed slowing down of the mitotic waves during consecutive division rounds. Considering that we experimentally observed a 2-4% longer period at the margin, close to the theoretically predicted 1.9%, this suggests that a linear gradient of cell cycle periods, rather than cell-cell coupling, gives rise to the observed radial mitotic waves found during embryonic cleavages.

### Increased cell-cell coupling reshapes the mitotic wave and prevents desynchronization

Next, to challenge our conclusion that the mitotic waves are produced in a coupling-independent manner, we asked how they would appear if the cell cycles were instead coupled. To that end, we performed numerical simulations of the Kuramoto model of interacting oscillators on a hemisphere with free boundary conditions^20^ (Fig. 2A). The oscillators were placed on a Fibonacci lattice covering the hemisphere, with nearest neighbors obtained with Delaunay triangulation. The phases of the oscillators obeyed the following equation: 𝛿_𝑡_𝜃_𝑖_ = 𝜔_𝑖_ + 𝜖 ∑_𝑗∈𝑁𝑖_ 𝑠𝑖𝑛(𝜃_𝑗_ − 𝜃_𝑖_) where 𝜔_𝑖_ represents the natural frequency of the *i*-th oscillator, and the second term describes the interactions between neighboring oscillators parametrized by the interaction strength 𝜖, with *Ni* being the nearest neighbors of the *i*-th oscillator. Due to cell-to-cell variations, we further assumed the presence of noise in the cell cycle lengths. Mathematically, we decomposed the frequencies into two parts 𝜔_𝑖_ = 𝜛(𝑧) + 𝛬_𝑖_, where the first term 𝜛(𝑧), is the inherent AP-to-margin oscillation gradient, and the second term is an independent spatial noise with covariance ⟨𝛬_𝑖_𝛬_𝑗_⟩ = 𝜎𝛿_𝑖𝑗_. Simulations with small interaction strength (ɛ) and low noise (σ) recapitulated the wild-type behavior, where mitotic waves originated at the AP, and accumulated delays causing a gradual slowdown over time (Fig. 2B-C). In contrast, the introduction of greater interaction strength and noise led the system to self-organize into smooth mitotic waves originating at the margin, similar to the case in *Xenopus* egg extracts, even though the ‘natural’ cell cycle lengths at the AP were preset to be shorter on average^21^ (Fig. 2B, D). For even larger values of noise, multiple waves emerged from random points of the simulated embryo (Fig. 2B). Overall, this theoretical phase diagram suggests that wild-type embryos are characterized by low to intermediate noise and very little coupling, and that increasing cell coupling should, in theory, shift the origin of the mitotic wave from the AP to the blastoderm margin.

**Fig. 2:**
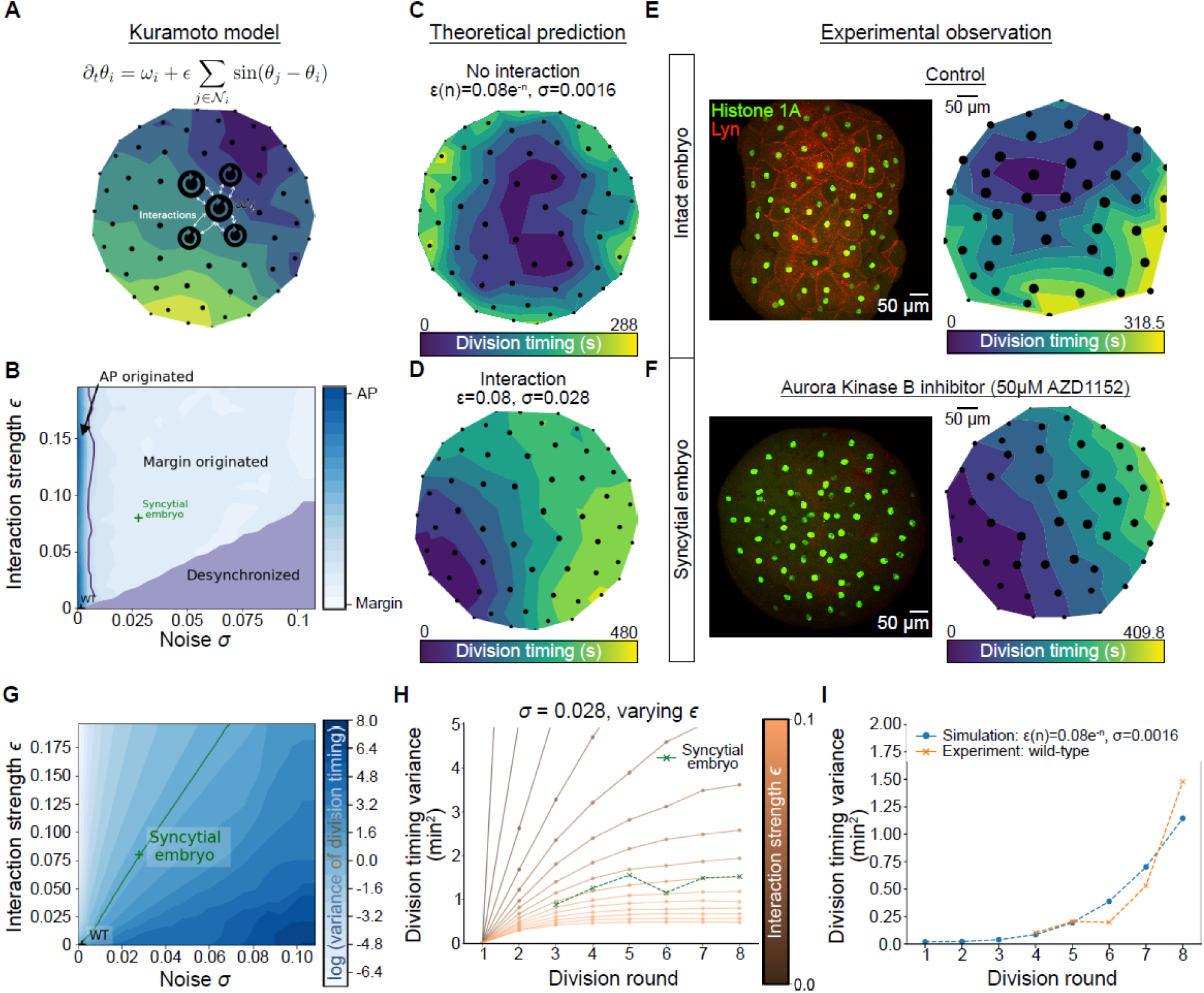
Cell-cell coupling repositions the mitotic wave origin to the margin and prevents desynchronization. (A) Schematic representation of our adapted Kuramoto model and the equation describing the evolution of the phase of a cell cycle. Equation: *θ_i_* represents the cell cycle phase of cell *i*, and *N_i_* denotes its nearest neighboring cells, determined through Delaunay triangulation of the hemisphere. The coupling strength *ɛ* represents how strongly each cell influences its neighbors’ timing. The intrinsic frequency *ω_i_* represents how quickly cell *i* would progress through its cell cycle in isolation. Model: Each black dot represents a cell positioned on a hemispherical Fibonacci lattice. Dot size increases with height (*z*) to aid visualization in this top-down projection. Lines between dots (not shown) indicate coupled neighbors that can influence each other’s cell cycle timing. (B) Phase diagram showing origin of the waves in *ε*-*σ* space. The boundary between AP-originated and margin-originated is thresholded at *z*=0.5, and the region of desynchronization corresponds to ⟨*e^iθ^j*⟩j < 0.6, where ⟨⟩j denotes averaging over all sites. (C) Contour plot of the radial mitotic wave across all surface cells at the 8th cleavage division round as predicted using the Kuramoto model of interacting oscillators, assuming an AP-margin period gradient of 1.9% and exponentially decaying cell-cell interaction strength (ε=0.08e^-n^, *σ*=0.0016). (D) Contour plot of the mitotic wave across all surface cells at the 8th cleavage division round as predicted using the Kuramoto model of interacting oscillators, assuming an AP-margin period gradient of 1.9% and partial interaction between cells (*ε*=0.08, *σ*=0.028). (E) Left: Representative image of a 128-cell stage control Tg(*actb2:lyn-tdtomato*) embryo injected with 1pg Histone1-Alexa Fluor 488 at the one-cell stage to show the presence of plasma membrane between individual nuclei. Right: Contour plot of the radial mitotic wave across all surface cells at the 8^th^ cleavage division round observed in a control embryo, where nuclear divisions were visualized using either Histone1-Alexa Fluor 488 injections or using Tg(*actb2:rfp-pcna*) embryos. n=7 embryos. (F) Left: Representative image of a 128-cell stage Tg(*actb2:lyn-tdtomato*) embryo injected with 1pg Histone1-Alexa Fluor 488 at the one-cell stage and treated with 50μM AZD1152 to show the absence of plasma membrane between individual nuclei. Right: Contour plot of the mitotic wave across all surface cells at the 8^th^ cleavage division round observed in a representative syncytial embryo, where nuclear divisions were visualized using either Histone1-Alexa Fluor 488 injections or using Tg(*actb2:rfp-pcna*) embryos. n=7 embryos. (G) Phase diagram of the Kuramoto model in *ε*-*σ* space. The log of the variance of the division timings at the 8^th^ round is plotted. The green solid line indicates the contour line of the observed variance at the 8^th^ round in syncytial embryos. (H) Variances of division timings for each round for fixed noise value, *σ*=0.028 as *ε* varies. Green dotted curve shows the variances measured in syncytial embryos. (I) Variances of division timings for each round in wild-type embryos (orange) fitted with a curve described by exponentially decaying interaction strength, *ε*(*n*)*=*0.08*e^-n^*, and low noise, *σ*=0.0016 (blue).

To experimentally test this prediction, we syncytialized cleavage-stage embryos by preventing cytokinesis - while allowing karyokinesis - through transient inactivation of Aurora Kinase B^22^ (Fig. 2E–F). We reasoned that such a syncytialized embryo would resemble conditions observed in *Drosophila* embryos and *Xenopus* egg extracts - both syncytial systems in which cell cycles are partially or fully synchronized via coupling-dependent trigger waves - and would therefore function as a coupled system. Remarkably, consistent with our theoretical predictions, we found that in syncytial embryos, the mitotic wave originated at the margin rather than at the AP, as seen in wild type embryos (Fig. 2B, D, F; Movies S1, S4). Moreover, by analyzing the wave characteristics at the 8th division round in syncytial embryos, we were able to estimate the effective ratio of coupling and noise, two competing factors promoting synchronization and desynchronization, respectively (Fig. 2G).

Notably, our model also predicted key differences in the evolution of cell cycle synchronization - measured by the variance in division timing across cleavage rounds - between wild type and syncytial embryos: in the absence of coupling (wild type embryos), noise is expected to accumulate progressively, resulting in a monotonically increasing variance in division timing. In contrast, in the presence of coupling (syncytial embryos), this variance should plateau over time, with a characteristic time scale determined by the coupling strength (Fig. 2H). Strikingly, our experimental observations confirmed this prediction. In syncytial embryos - unlike in wild types - ‘cells’ continued to cycle metasynchronously even beyond the 10th division, indicating strong coupling between cell cycles (Fig. S3A; Movies S1, S4). The temporal evolution of variance closely matched our theoretical prediction, particularly for ε = 0.08 and σ = 0.028 (Fig. 2H). In contrast, wild type embryos exhibited continuously increasing variance, with an even faster temporal scaling than predicted by the model during later cleavages (Fig. 2I). Interestingly, this discrepancy could be quantitatively explained by the incomplete nature of cytokinesis during early cleavages - suggesting the presence of transient coupling that rapidly diminishes as development progresses^23^. This interpretation is supported by both our experimental data and theoretical framework (see SI Theory Note for detailed assumptions and modeling approach).

Finally, to further challenge the notion of increased coupling of ‘cells’ in syncytial embryos, we injected *chek1 m*RNA into embryos before syncytialization and assessed whether they could resist premature desynchronization typically observed when *chek1* was overexpressed in wild type embryos (Figs. 1F, S1, Movie S3). We reasoned theoretically that the coupling inferred for syncytial embryos is large enough that it can override a significant increase in noise, such as the desynchronizing effects of *chek1* expression (Fig. 2B). Indeed, we found that in *chek1*-overexpressing embryos, the mitotic waves originated at the margin, and ‘cells’ continued to cycle metasynchronously even beyond the 10th division round, suggesting that cells in the syncytial embryo are, in fact, tightly coupled. (Fig. S3B-C, Movie S5). By extension, this implies that such a coupling is absent/minimal in wild type embryos and, consequently, that mitotic waves are produced largely independently of cell-cell interactions in wild-type embryos. Collectively, these findings suggest that cell cycles are only weakly coupled during cleavages and that metasynchrony arises predominantly by cell-autonomous processes.

### Early embryo geometry establishes a cell volume gradient along the AP-margin axis through guided asymmetric cell divisions

For the cell cycle to slow down from the AP to the margin in a predominantly cell-autonomous manner, there must be a factor causing unequal lengthening of the S-phase along the AP-margin axis. Among the most important factors regulating S-phase length is the nucleocytoplasmic (N/C) ratio^24,25^. A high N/C ratio causes S-phase lengthening, for instance, by retarding Cdk1 activation^24^. Consistent with such an effect of the N/C ratio on cell cycle length, we found that the length of the S-phase across the metasynchronous cleavage rounds negatively correlated with cell volume (Fig. 4A-B). Therefore, we hypothesized that an AP-margin gradient of cell volumes in the cleavage-stage embryo might be responsible for unequally lengthening cell cycles from the AP to the margin. To test this hypothesis, we measured cell volumes by semi-automatically segmenting surface cells at the 128-cell stage (round 8). Indeed, we found that, on average, cells closer to the AP were significantly larger than those away from it (Figs. 3A, 4G). To determine at what stage such a volume gradient first becomes apparent, we also measured cell volumes in embryos at the 8-cell stage (round 4) and the 16-cell stage (round 5) - the first stages where cells can be classified as being ‘central’ or ‘peripheral’ based on their position. Interestingly, we observed that cells closer to the AP were consistently larger than the marginal cells at both these stages, showing that the AP-margin gradient of cell volumes arises already in the very early embryo (Fig. 3B, C). However, no such gradient was observed for the nuclear volume, indicating that the peripheral cells have a larger N/C ratio (Fig. 3B). Furthermore, the cell volume gradient not only persisted but was also further enhanced by subsequent cell divisions, as mother cells having unequal sizes themselves again divided asymmetrically to produce yet unequally sized daughters, thereby further diversifying cell volumes through cell lineage in addition to the already established position-driven cell volume diversity (Fig. 3D). For such unequal cell divisions to robustly give rise to a gradient in cell volumes along the AP-margin axis, upon division the daughter cell closer to the AP must be on average larger than the daughter cell further from the AP. In line with this, we found that cell divisions in the early cleavage stages showed a tendency to produce larger daughter cells closer to AP, with the central daughter cell (closer to AP) being, on average, ∼12.4% larger by volume than its peripheral sister cell (away from AP) (Fig. 3E).

**Fig. 3:**
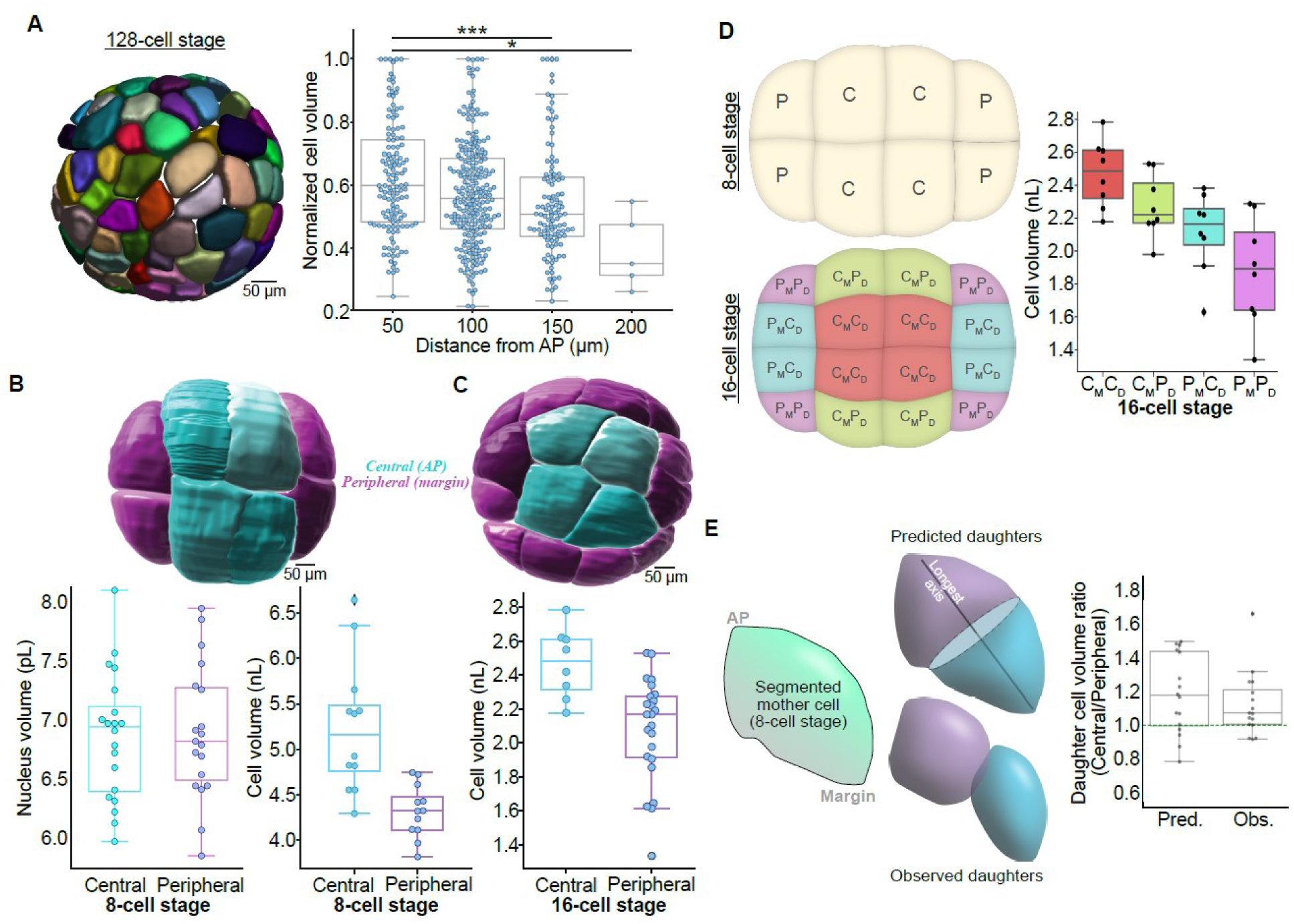
Embryo geometry drives asymmetric cell divisions to generate a gradient of cell volumes in the early embryo. (A) Representative image of a 128-cell stage embryo (round 8) with surface cells segmented (left) and box-plot showing normalized cell volumes as a function of cell position with respect to the AP (right). Cell volumes were internally normalized by dividing each cell volume by the volume of the largest cell in the embryo. Each dot represents the data for an individual cell. (n=469 cells, 12 embryos). (B) Representative image of an 8-cell stage embryo (round 4) with surface cells segmented (top) and box-plot showing volumes of individual nuclei and cells based on their position relative to the AP (bottom). ‘Central’ cells are positioned at AP, whereas ‘peripheral’ cells are positioned away from the AP (n=20 central and 19 peripheral nuclei, 5 embryos; 12 central and 12 peripheral cells, 3 embryos). (C) Representative image of a 16-cell stage embryo (round 5) with surface cells segmented (top) and box-plot showing volumes of individual cells based on their position relative to the AP (bottom; n=8 central and 24 peripheral cells, 2 embryos). (D) Schematic showing the lineage relationship between cells at the 8-cell stage (top-left) and the 16-cell stage (bottom-left). At the 8-cell stage, cells marked ‘C’ are centrally positioned, and those marked ‘P’ are peripherally positioned. At the 16-cell stage, cells are labelled to indicate both their positions (CD, centrally; PD, peripherally) and the identities of their mother cells (CM, centrally positioned mother cell; PM, peripherally positioned mother cell). Box plot showing the distribution of cell volumes at the 16-cell stage based on each cell’s position and lineage identity (right) (n=8 cells for each category, 2 embryos). (E) Schematic showing the pipeline for predicting daughter cell volumes at the 16-cell stage (left). Each segmented cell surface from the 8-cell stage was used to identify the longest cell axis, which was then bisected to obtain daughter cells at the 16-cell stage. Violin plot showing a comparison between the ratios of volumes of the central and peripheral daughter cells obtained from each cell division, respectively, either predicted (Pred.) or experimentally measured (Obs.) (right; n=16 cell pairs, 2 embryos).

**Fig. 4:**
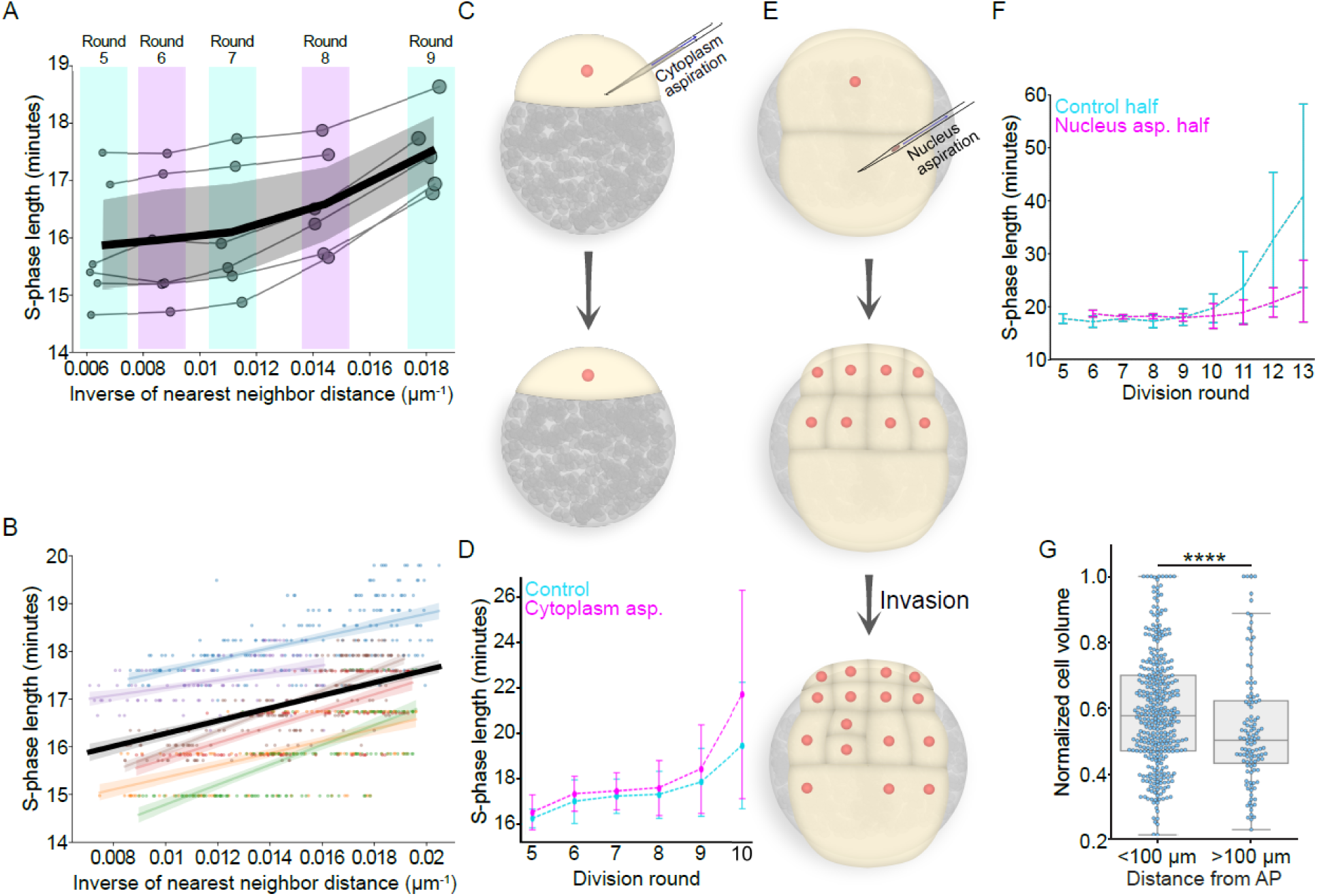
Cell volume regulates S-phase length (A) Line plots showing mean S-phase lengths in individual (thin, grey lines) and across (thick, black line, mean ± 95% CI) embryos as a function of the inverse of the mean nearest neighbor distance (mean of 3) in the metasynchronous cell cycle stages. Division rounds are indicated by the size of the dots (smallest: division round 5, largest: division round 9 (n=6 embryos) and also indicated at the top. (B) Linear regression plots showing the relationship between the inverse of the nearest neighbor distance (mean of 3) for each cell and the length of its S-phase. Dots represent single cells, whereas the lines show the best-fit linear curve for the data. Each color represents the data for a unique embryo. The thick, black line shows a summary best-fit for the data of all cells across all embryos. For each embryo, cells only within the 5^th^-95^th^ percentile of their nearest neighbor distances and S-phase length were considered for the analysis, which minimizes erroneous measurements at the extremes but also underestimates the relationship (n=6 embryos, 964 cells). (C) Schematic representation of the experiment where a fraction of the cytoplasm was aspirated from the blastodisc within approximately 10 minutes before the first cell division to increase the nucleocytoplasmic ratio. (D) Representative line plot showing S-phase length in each division round in control embryos and embryos from which cytoplasm was aspirated (mean ± SD, n≥3 embryos). (E) Schematic representation of the experiment where one of the nuclei at the two-cell stage was aspirated to decrease the nucleocytoplasmic ratio in the aspirated half of the embryo (top). Only the nucleated, control half initially develops (i.e., shows mitotic cycles) (middle) until nuclei from this half invade the enucleated half (bottom), which then begins to develop with a smaller nucleocytoplasmic ratio. (F) Representative line plot showing S-phase length in each division round in the control and enucleated halves of the embryo depicted in (E) (mean ± SD, n=3 embryos). (G) Box plot showing normalized cell volumes as a function of cell position with respect to the AP. Cell volumes were internally normalized by dividing each cell volume by the volume of the largest cell in the embryo. Each dot represents the data for an individual cell. Data is the same as in Fig. 3A, but the cells here have been classified into only two bins (n=469 cells, 12 embryos; Student’s t-test, p<0.001).

To determine what causes such patterned asymmetric cell divisions, we investigated how cell divisions are oriented and the cleavage plane positioned in the early embryo. While several sophisticated models have been proposed to explain the orientation and positioning of the cleavage plane across systems, we started by first testing if following the simplest model, the ∼140-year old Hertwig’s rule, can recapitulate the experimentally observed pattern of asymmetry^26,27^. More specifically, we assumed that the mitotic spindle in dividing blastomeres would orient along the longest cell axis, thereby dividing the cell along this axis. To test if such a geometry-driven mechanism would suffice to generate the observed pattern of asymmetric cell divisions and, therefore, the cell volume gradient, we predicted daughter cell volumes at the 16-cell stage by first identifying the longest axes of segmented cells at the 8-cell stage through Principal Component Analysis and then bisecting them to obtain two daughter cells from each division^28^ (Fig. 3E). Indeed, we found that with this approach, the predicted daughter cell size asymmetry closely matched the observed asymmetry for most cell divisions, such that the central daughter cells (closer to the AP) at the 16-cell stage were, on average, ∼18.5% larger than their peripheral sister cells (further away from the AP) by volume (Fig. 3E). Importantly, this tendency of central daughter cells being bigger than peripheral ones is determined by the specific geometry of the first cell of the fertilized egg, the blastodisc, that is delineated not only by a plasma membrane at its outer side, but also by the interface to the yolk granules on its inner side^10,29^. As the interface to the yolk granules is curved, the first two cell divisions will give rise to equally sized daughter cells that are more pointed and narrow at their peripheral ends. Dividing these cells again along their long axis will then, as a result, generate daughter cells where the central cell is larger than the peripheral one. This suggests that a gradient of cell volumes along the AP-margin axis in the cleaving embryo can be explained by the specific geometry of the first cell in the fertilized egg, leading to unequal cell division along the AP-margin axis during subsequent division rounds.

### Cell volume gradient causes patterned cell cycle lengthening along the AP-margin axis

Whereas the influence of cell volume on cell cycle length has been established in several systems, its role in zebrafish embryos remains unclear due to conflicting reports regarding the determinative effect of cell size in this context^13,15^. To unequivocally assess the contribution of cell volume - and, by extension, the nucleocytoplasmic (N/C) ratio - to the regulation of cell cycle length, we first increased the N/C ratio at the one-cell stage by aspirating cytoplasm from the blastodisc (Fig. 4C). Consistent with the known role of elevated N/C ratios in prolonging cell cycles, we observed that, across different embryos, the S-phase either lengthened prematurely or showed greater lengthening during the 10th division round, and the cell cycles desynchronized earlier than in control embryos (Fig. 4D). To test the converse, we decreased the N/C ratio by replicating an experiment originally conducted by Kane and Kimmel, aspirating one nucleus from embryos at the two- or four-cell stage^13^ (Fig. 4E). In these embryos, only the control half - containing the nucleus - underwent initial cleavages, while no divisions occurred in the enucleated half (Fig. 4E; Movie S6). However, in a subset of embryos, nuclei from the control half eventually invaded the enucleated half around the 4th or 5th division, prompting the onset of cleavages in that region (Fig. 4E; Movie S6). Notably, although all nuclei had undergone the same number of division cycles by this stage, those in the previously enucleated half now resided in a substantially larger cytoplasmic volume - comparable to the blastomere volume of cells three to four divisions earlier - and therefore had a lower N/C ratio. Consistent with earlier findings, these nuclei did not exhibit S-phase lengthening during the 10th division round - the point at which cell cycles typically begin to desynchronize in the control half - but instead continued to cycle metasynchronously. This supports the notion that a higher N/C ratio promotes S-phase lengthening and contributes to the onset of cell cycle desynchronization beyond the 10th division (Fig. 4F).

Having confirmed that the N/C ratio, in principle, can influence the S-phase length, we asked how perturbing the embryo-scale cell volume gradient within the cleaving embryo would affect cell cycle (meta)synchrony and, consequently, the mitotic wave. To alter the cell volume gradient, we first generated bilobed embryos, displaying two ectopic (morphological) ‘animal poles’, by mounting two-cell stage embryos in low melting point agarose (0.67% w/v in E3 buffer) when the first cytokinesis was in progress (Fig. 5A). We reasoned that spatially confining and, therefore, reshaping the embryo in this manner would produce larger cells away from the AP at the two ectopic animal poles. Interestingly, we found that not only the gradient of cell volumes was reshaped in the embryo, but also that two mitotic waves emerged, one from each of the two ectopic APs, supporting the notion that the cell volume gradient is linked to the emergence of the mitotic wave (Fig. 5A-B, Movie S7). To further challenge this suggestion, we removed a part of the yolk on the vegetal side of the two-cell stage embryo to increase the curvature of the blastodisc (the two blastomeres positioned on the yolk cell; Fig. 5C). Based on our analysis indicating that the cell volume gradient is a product of the curvature of the blastodisc-yolk interface, we reasoned that this increased curvature would reorient the longest axes of the two cells, such that they align more along the animal-vegetal (AV) axis, which would then amplify the volume asymmetry of cleavage divisions and, consequently, the phase difference between cell cycles (Fig. 5C). Strikingly, we found that in yolk-severed embryos, with cell divisions more aligned along the AV axis, the mitotic wave indeed showed a greater cell division timing variance compared to the control embryos (Fig. 5C-D, Movie S8). Taken together, these observations strongly support the idea that a gradient of cell volume along the AP-margin axis, guided by embryo geometry, causes the generation of a gradient of S-phase lengths along the AP-margin axis of the blastoderm, thereby generating radial mitotic waves in the zebrafish embryo.

**Fig. 5:**
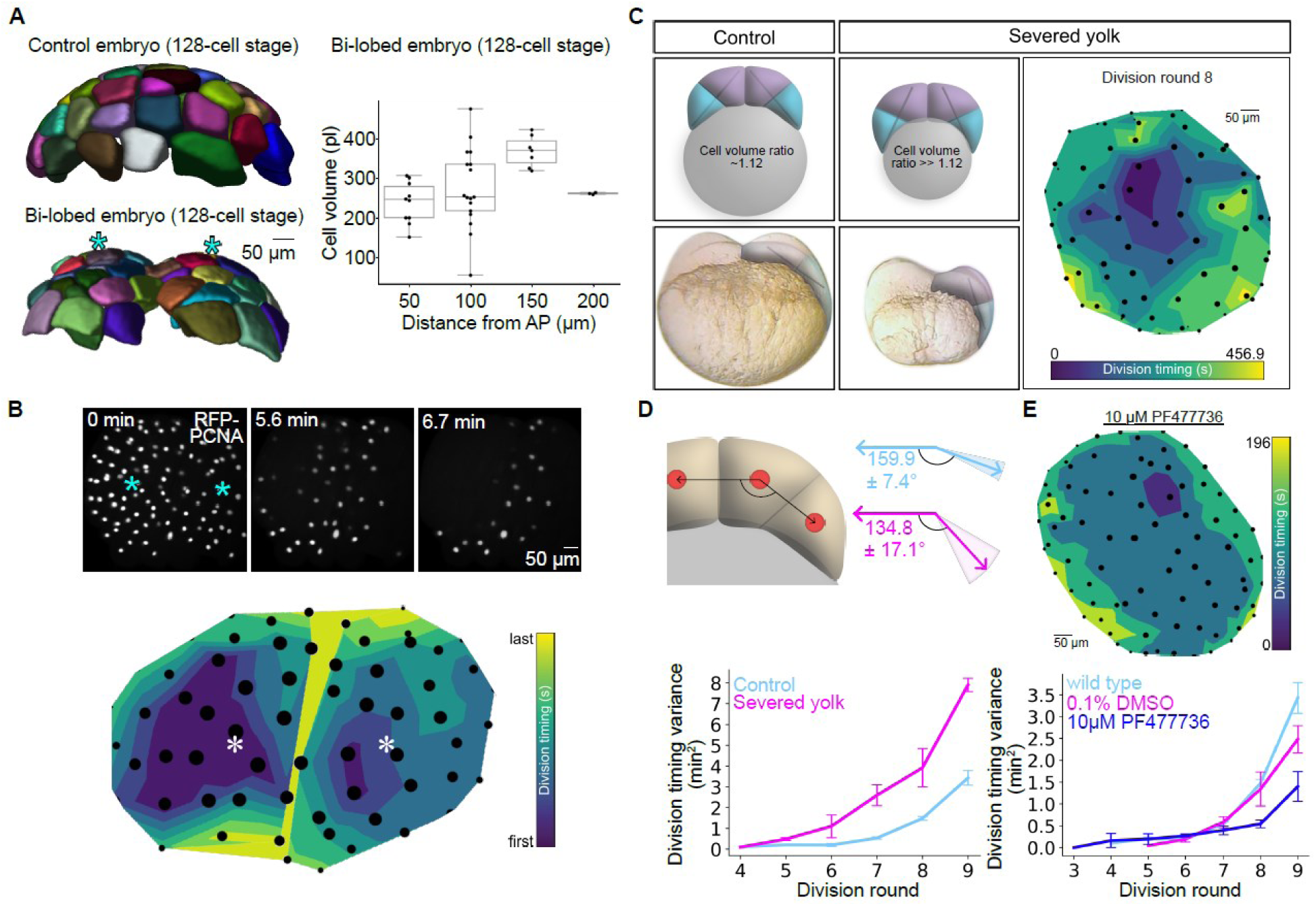
AP-margin cell volume gradient increases cell cycle metasynchrony in a Chek1-dependent manner (A) Representative images of 128-cell stage (round 8) control and bilobed embryos with surface cells segmented (left) and box-plot showing cell volumes as a function of cell position with respect to the AP (right) for this bilobed embryo. Cyan asterisks mark the ectopic animal poles in the bilobed embryo. (B) Montage of images (top) and a contour plot (bottom) showing the propagation of mitotic waves in a representative bilobed Tg(*actb2:rfp-pcna*) embryo at the 8^th^ division round. Asterisks mark the ectopic animal poles in the bilobed embryo (n=5 embryos). (C) Schematic representations (top-left) and representative images (bottom-left) of a control embryo and an embryo where the yolk has been severed at the two-cell stage to increase the curvature of the blastoderm-to-yolk cell interface. Lines inside the cells indicate the longest cell axis along which the divisions (division round 3) are expected to occur. Cells marked in magenta and cyan represent the resulting central and marginal daughter cells, respectively. Contour plot (right) shows mitotic wave propagation in the 8^th^ division round of a yolk-severed embryo (AP-view). (D) Schematic representation (top) of the decreased angle of cell divisions in the 4^th^ division round in yolk-severed embryos due to an increased curvature of the blastoderm-to-yolk cell interface compared to corresponding control embryos (angles are mean ± SD; n>10 cells). Line plot (bottom) showing the variance in cell division timings in control and yolk-severed embryos (mean ± SD; n=4 embryos). (E) Representative contour plot (top) showing mitotic wave propagation in a Chek1 inhibitor (PF477736)-treated embryo in division round 8 (AP-view). Line plot (bottom) showing the variance in cell division timings in control and PF477736-treated embryos (mean ± SD; n=4 embryos).

Among the major responders to a high N/C ratio is Chek1, which, when active, delays the S-M transition^24^. Therefore, having determined that a gradient of cell volume patterns cell cycle length in the zebrafish embryo, we inquired whether Chek1 mediates the effect of cell volume on S-phase length. To that end, we inhibited Chek1 activity by treating embryos with 10 µM PF477736, a specific Chek1 activity inhibitor^30^. If Chek1 activity is required for the gradual metasynchronization of cell cycles through unequal S-phase lengthening, the cell cycle progression would be expected to show lesser variation in the treated embryos. Consistent with this, we found that, while Chek1-inhibited embryos produced mitotic waves like those in control embryos, they indeed exhibited a lower mitotic division timing variance (Fig. 5E, Movie S9). This suggests that Chek1 activity is critical for translating a cell volume gradient into a mitotic wave along the AP-margin axis of the blastoderm.

### Cell volume/cell cycle period gradients pattern ZGA and germ layer specification along the AP-margin axis

Finally, we asked whether the geometry-derived cell volume gradient - and the resulting patterned mitotic metasynchrony - plays a developmental role that extends beyond the cleavage stages and impacts later embryogenesis. In zebrafish embryos, mitotic metasynchrony diminishes in parallel with the onset of zygotic genome activation (ZGA), a pivotal process that lays the foundation for cell fate heterogeneity and is central to metazoan development^13^. Given previously reported roles of cell volume (N/C ratio promotes ZGA) and cell cycle length (shorter cell cycle prevents ZGA) in regulating ZGA, we investigated whether the observed gradient in cell volume and mitotic metasynchrony influences the onset of zygotic transcription^31–36^. To address this, we tracked the timing and spatial pattern of zygotic transcription initiation by imaging endogenous *pri-miR430* transcripts using MOrpholino VIsualisation of Expression (MOVIE)^37^ (Fig. 6A). *miR430* genes are among the earliest and most abundantly transcribed genes during zebrafish ZGA, making them ideal markers for assessing transcriptional onset^38,39^.

**Fig. 6:**
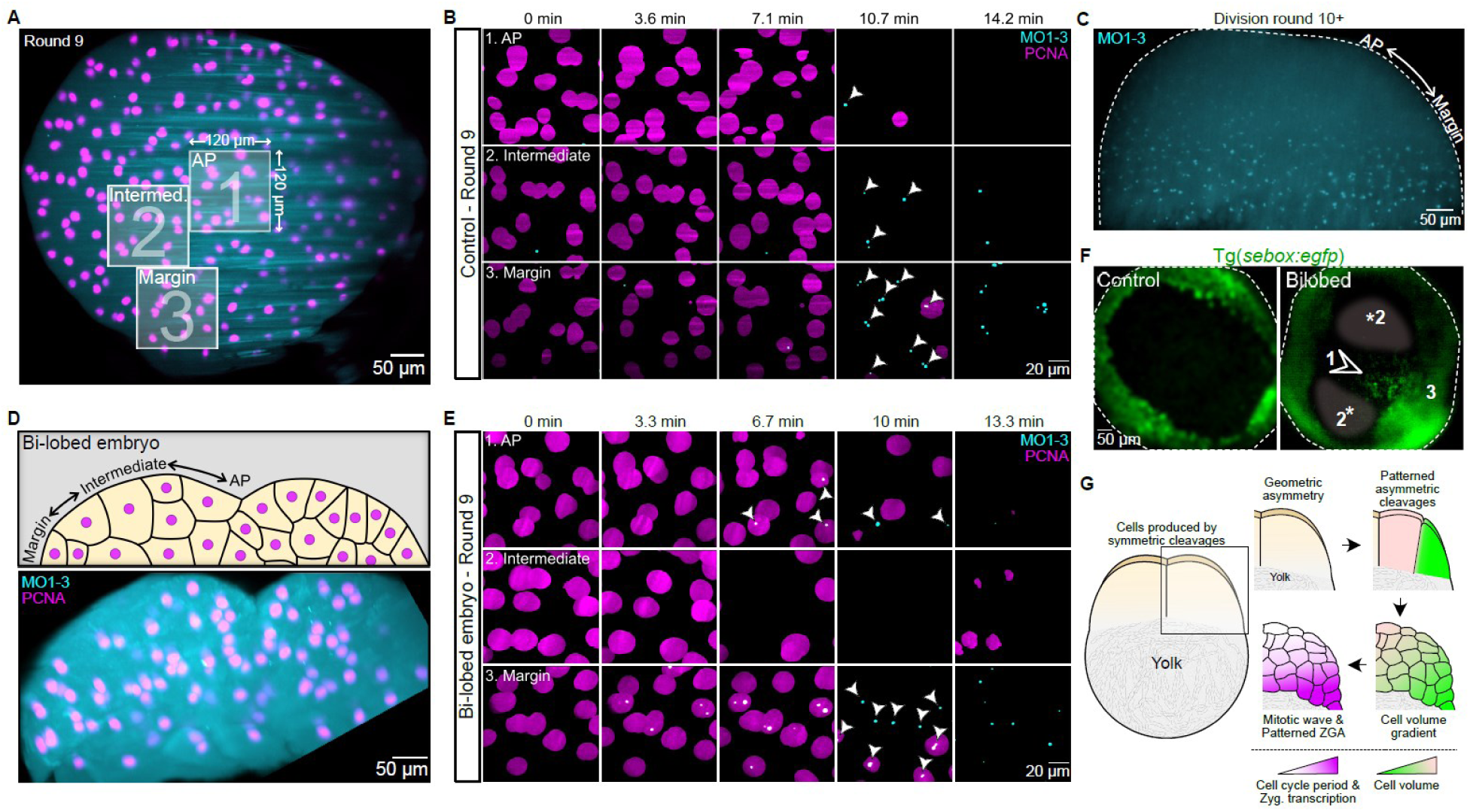
Zygotic transcription is initiated in a patterned manner along the AP-margin axis (A) Representative maximum intensity projection image showing the animal pole-view of a control Tg(*actb2:rfp-pcna)* embryo injected with MO1-3-Fluorescein in the 9^th^ division round. Cyan, MO1-3-Fluorescein; magenta, nuclei (PCNA). The three insets, each 120 µm X 120 µm in size, demarcate the regions (1, animal pole; 2, intermediate region; 3, margin) whose higher-magnification images are shown in (B) and (E). (B) Zoomed-in views of the insets shown in A. Cyan dots represent nascent *mi430* transcripts detected by MO1-3-Fluorescein (n=4 embryos). (C) Lateral view of a control embryo in the division round 10 injected with MO1-3-Fluorescein. (D) A schematic (top) and a representative orthogonal maximum intensity projection image (bottom) of a bi-lobed Tg(*actb2:rfp-pcna)* embryo injected with MO1-3-Fluorescein embryo in the 9^th^ division round. Cyan, MO1-3-Fluorescein; magenta, nuclei (PCNA). (E) Zoomed-in views of the equivalent regions (1, animal pole; 2, intermediate region; 3, margin) demarcated in (A) for a bi-lobed embryo in the 9^th^ division round. Cyan dots represent nascent *mi430* transcripts (n=3 embryos). (F) AP-view of control and bilobed Tg(*sebox:egfp*) embryos, with comparable regions from (A) indicated in the bilobed embryo. Arrowhead indicates the presence of ectopic *sebox:egfp* expression at the original AP, where a valley is formed between two domes (ectopic APs) of the bilobed embryo (n=16 embryos). (G) Schematic model representing geometry-driven asymmetric cell division patterning in the early embryo. Patterned asymmetric divisions generate a gradient of cell volume, producing mitotic phase waves and spatially patterning ZGA onset along the AP-margin axis.

In most embryos, we observed the first signs of *miR430* transcription during the 9th division round. Notably, during rounds 9 and 10, transcription was initiated in a graded manner: *miR430* transcription foci first appeared - and/or persisted longer - in cells near the blastoderm margin. They then emerged in an intermediate zone between the margin and the AP, and finally in cells located at the AP (Figs. 6B-C, S4; Movies S10, S11). These findings suggest that zygotic transcription begins as a spatiotemporal gradient along the AP-margin axis, with marginal cells exhibiting higher transcriptional activity or competence than AP cells over the course of one to two division rounds. To determine whether this pattern of ZGA onset is driven by the underlying gradients in cell volume and/or cell cycle length, we examined bilobed embryos in which both of these gradients are perturbed (Figs. 5A–B, 6D). In contrast to control embryos, cells at the AP in bilobed embryos are smaller and cycle more slowly than those in the intermediate zone (Figs. 5A–B). Remarkably, in these embryos, *miR430* transcription was initiated first at both the margin and the AP - where cells are smallest - before appearing in the intermediate region (Fig. 6E; Movie S12). These results suggest that the gradient of cell volume and/or cell cycle length along the AP-margin axis influences the spatiotemporal onset of zygotic transcription. Thus, early embryo geometry can be linked directly to the initiation of zygotic gene expression, providing a mechanistic bridge between physical form and transcriptional timing in early development.

To determine whether the geometry-derived transcriptional gradient within the embryo plays a role in embryonic development, we asked whether altering the spatiotemporal pattern of ZGA onset in bilobed embryos would affect the specification of the first cell fates within the blastoderm. To investigate this, we used *Tg(sebox:egfp)* embryos to visualize the earliest mesendoderm progenitors, specified at the onset of gastrulation^40,41^. In control embryos, mesendoderm progenitors were exclusively specified at the blastoderm margin - the region where ZGA is first initiated. In contrast, approximately 30% of bilobed embryos exhibited mesendoderm progenitors not only at the margin but also ectopically within the ‘valley’ between the two domes, reflecting the altered ZGA pattern in these embryos (Fig. 6F). These findings suggest that zygote geometry triggers a cascade of developmental events necessary for correct cell fate specification - an essential aspect of ensuring developmental robustness (Fig. 6G).

## Discussion

Previous studies have shown that the shape of the zebrafish egg - particularly the blastodisc, the region from which all embryonic cells and tissues originate - is established by ooplasmic streaming, which segregates yolk granules from the cytoplasm within the fertilized egg^29,42^. Building on this, our findings suggest that the initial shape of the blastodisc drives asymmetric cell divisions during successive reductive cleavages by guiding mitotic spindle orientation, based on the simple premise that the spindle consistently aligns with the cell’s longest axis, as described by Hertwig’s rule^27,43,44^. These asymmetric divisions progressively generate a gradient of cell volumes along the AP-margin axis of the developing blastoderm, which, in turn, influences both cell cycle progression and ZGA along the same axis. While we do not rule out the potential contribution of more complex mechanisms linking blastodisc shape to asymmetric cell divisions - such as differential interactions between spindle microtubules and yolk granules, the cell cortex, or the plasma membrane - our results suggest that such factors are not required to explain the emergence of asymmetric divisions. Thus, the formation of a gradient in cell volume, and the consequent regulation of cell cycle dynamics and ZGA patterns, appears to be a direct outcome of the blastodisc’s geometry.

Interestingly, our data connect the observed gradients in cell volume and/or cell cycle length to the onset of ZGA. This aligns with recent findings in *Xenopus* embryos, where ZGA is first initiated in the smaller cells at the animal pole^13^. Multiple molecular mechanisms have been proposed to explain the link between cell cycle length and the onset of ZGA, including the titration of cytoplasmic transcriptional repressors as the nucleocytoplasmic (N/C) ratio increases, and/or progressively lengthening cell cycles that permit sustained zygotic transcription. Our observation that ZGA not only initiates earlier but also persists longer in the smaller, marginal cells suggests that both mechanisms may contribute to the regulation of zygotic gene activation.

We have previously shown that radially patterned cell behaviors and asymmetric tissue compaction along the AP-margin axis of the blastoderm emerge at the onset of gastrulation, providing essential conditions for proper morphogenetic movements during this stage of development^19,45^. It is plausible that these differences in tissue properties originate from distinct transcriptional profiles and dynamics, which are themselves driven by the differential patterns of ZGA described in this study.

Our findings thus establish a likely mechanistic link between two critical developmental milestones: the determination of blastodisc geometry immediately after fertilization and the onset of asymmetric tissue fluidization during gastrulation. This connection is further supported by our earlier observation that asymmetric tissue fluidization depends on proper mesendoderm specification^19,45^, and by our current finding that embryos with perturbed geometry exhibit mesendodermal fate misspecification that parallels the altered pattern of ZGA onset. This suggests that inherent geometric asymmetries in blastodisc shape can have long-lasting developmental consequences, manifesting only later as their effects accumulate during the early proliferative phase of embryogenesis.

Consistent with this notion, several other organisms - including *C. elegans*, sea urchins, and *Xenopus* - exhibit alignment between axes of cell cycle length and/or cell volume and the axes along which germ layer fates diverge, underscoring the developmental significance of geometry-derived patterns as robust and reproducible determinants of embryonic organization. Nonetheless, the specific outcomes of this fundamental relationship are likely to vary across species and must be examined within the unique developmental context of each organism.

## Methods

### Cell cycle analysis

Tg(*actb2:rfp-pcna*) embryos were dechorionated and, at the desired stage, mounted in 0.5% agarose gels prepared in E3 buffer with the animal pole facing upward in casts made with 2% agarose solution in E3 buffer^12^. RFP fluorescence was imaged at 23.5 °C using a 20x water-dipping objective on Zeiss LSM800 (N.A.=0.8), LSM880 (N.A.=1.0), or LSM900 (N.A.=1.0) upright confocal microscopes. 200-350 µm-thick Z-stacks were acquired for different experiments with temporal resolutions ranging between 17-65 sec/Z-stack. The presence or absence of nuclear RFP-PCNA fluorescence was used to determine if a cell was in the S or M phase of the cell cycle, respectively^46–48^. The nuclear signal was segmented in Fiji using the Labkit plugin^49,50^. The segmented objects were imported into Bitplane Imaris (https://imaris.oxinst.com/) to generate tracks and obtain tracking-related data, including track start time (S-phase entry), track end time (M-phase entry), track duration (S-phase length), and the nuclear coordinates at the last S-phase time point (position). These data were then analyzed using the Pandas Python library to calculate, for instance, the division timing variance in a given division round, followed by plotting using mainly the Matplotlib and Seaborn libraries^51–53^. Anaphase onset timings for comparison between sister and non-sister cells were determined by generating 3D image stacks of wild-type AB embryos injected with 0.5 pg Histone 1-Alexa Fluor 488 (Catalog #H13188, Invitrogen) at the one-cell stage as described above.

### Mitotic wave speed

The mitotic wave speed for a given round of division was determined by first identifying the position of the wave origin and the timing of the wave origination, measuring all other nuclear distances and mitotic entry timings relative to this position and time, and then generating the best-fitting distance-time linear curve. In the cases where more than one nucleus could be considered the origin of the mitotic wave due to a simultaneous entry into mitosis, the mean of the 3D coordinates of such nuclei was considered the position of the wave origin.

### Microinjections

To visualize cell cycle progression using Histone 1 as the marker, 0.5 pg Histone 1-Alexa Fluor 488 was injected at the one-cell stage into wild-type AB or Tg(*actb2:lyn-tdTomato*) embryos^54^. Mosaic perturbation of the cycling period of a subset of the cells was carried out by mounting 32/64-cell stage embryos in 2% agarose solution in E3 buffer and injecting 0.2-0.3 pg of Histone 1-Alexa Fluor 488 into such cells. *chek1* overexpression was achieved by injecting 12 pg *chek1* mRNA into the one-cell embryo^19,45^. mRNA was prepared as described previously^21^.

### Drug treatments

Aurkb function was inhibited by transiently incubating dechorionated <5 min post fertilization (mpf) embryos in 50 µM AZD1152 (Catalog #SML0268, Sigma-Aldrich) solution for 15 min, after which they were returned to E3 buffer. Embryos showing successful nuclear cycling despite a failure of cytokinesis were then imaged as described above. Inhibition of Aurkb function concomitantly with *chek1* overexpression or Histone 1-Alexa Fluor 488 visualization was performed by injecting 12 pg *chek1* mRNA or 0.5 pg Histone 1-Alexa Fluor 488, respectively, at the one-cell stage as described above, followed by incubation in the Aurkb inhibitor. Chek1 activity inhibition was performed by dechorionating <10 mpf embryos and incubating them in 10 µM PF477736 (Catalog #HY-10032, MedChem Express) solution in E3 buffer throughout the duration of analysis.

### Nuclear volume measurement

Nuclear RFP-PCNA fluorescence was segmented from Z-stacks by thresholding using Fiji, and their volumes were measured using the 3D Objects Counter plugin^55^.

### Cell volume measurement

Cell volumes at the 8- and 16-cell stages were measured by semi-automatically generating cell segmentation using the ‘Contour’ function for surface creation in Bitplane Imaris from Z-stacks of Tg(*actb2:mCherry-CAAX*) embryos^56^. Once segmented, the physical attributes of individual cells, such as volume and position, were exported from Bitplane Imaris as CSV files. The last time point before the onset of the fourth or fifth mitotic round was used to measure the cell volume for the 8- or the 16-cell stage, respectively. Cell volumes at the 128-cell stage were measured by generating cell segmentations using the Limeseg plugin in Fiji and exporting the cell volume data for individual segmented objects^57^. All downstream data analysis was performed in Python, mainly using the Pandas library, and the plots were produced using the Matplotlib and Seaborn libraries.

### Daughter cell volume prediction

Cell surfaces generated in Bitplane Imaris at the 8-cell stage were exported as WRL files, imported into MeshLab (https://www.meshlab.net/), from where individual cell surfaces were exported as PLY files. These files were subsequently processed in Python to identify the longest cell axis by performing Principal Component Analysis (PCA), along which the cell was bisected into two daughter cells representing cells at the 16-cell stage^28^.

### Nearest neighbor distance calculation

Nuclei positions were obtained from Bitplane Imaris, as detailed above. For each nucleus, the three nearest neighbors were identified, and their mean distance from the reference nucleus was calculated using the Pandas library in Python.

### Cytoplasm aspiration

Approximately 30 mpf Tg(*actb2:rfp-pcna*) embryos were mounted in E3 buffer on a substrate prepared with 2% agarose solution in E3 buffer. A 20 µm blunt-end needle attached to a 10 µl Hamilton syringe was inserted through the yolk into the blastodisc, and the cytoplasm was carefully aspirated. The nuclear RFP-PCNA signal was constantly observed using a Leica stereo-fluorescence microscope to ensure the nucleus was not extracted together with the cytoplasm.

### Nucleus aspiration

The setup and mounting procedure for the nucleus aspiration were the same as for cytoplasm aspiration described above, except that, in this case, the nucleus was aspirated rather than the cytoplasm and from embryos at the two-cell stage rather than the one-cell stage. Care was exercised to ensure that the nucleus was extracted with a negligible volume of the cytoplasm being aspirated in the process.

### Generation of bilobed embryos

Bilobed embryos were created by spatially confining dechorionated embryos at the two-cell stage, still undergoing cytokinesis from the first division round, in 0.5-0.6% agarose in E3 buffer. Microscopy was performed as described above.

### Yolk severing

To sever the yolk, dechorionated embryos at the two-cell stage were placed on a substrate of 3% methylcellulose in E3 buffer in a glass dish. Using an eyelash attached to the end of a Pasteur pipette, the vegetal half of the yolk was first punctured to prevent the embryo from exploding due to a buildup of internal pressure during the severing procedure, and then carefully sliced. The embryo was allowed to recover for at least 15 minutes after the procedure, followed by mounting and microscopy, as detailed above.

### Cell division angle measurement

Four orthogonal sections, each showing four nuclei along the 4th round division plane at the 16-cell stage, were generated using the Fiji Reslice function from Z-stacks of individual Tg(*actb2:rfp-pcna*) embryos. The angles between nuclei were then measured using Fiji, as represented in Fig. 5D, to obtain the angle of cell divisions in the 4th division round.

### miR430 transcription analysis

pri-*miR430* transcripts were labelled as described previously^37^. Tg(*actb2:rfp-pcna*) embryos injected with MO1-3-Fluorescein (Gene Tools) at the one-cell stage were mounted in 0.5% agarose solution in E3 buffer in #3 Fluorinated ethylene propylene (FEP) tubes, and Z-stacks were generated with a temporal resolution of approximately one min/Z-stack using a 20x/N.A. 1.33 objective on a Zeiss Lightsheet 7 microscope. Maximum intensity projections of these time-lapse recordings were then used to assess the transcription status of individual nuclei.

## Supporting information

Detailed description of the theoretical modelling and analysis of cell cycle synchronization.

## Acknowledgments

We thank Nicoletta Petridou (EMBL) for sharing results prior to publication. NM was supported by funding from the European Union’s Horizon 2020 programme under the Marie Skłodowska-Curie COFUND Actions ISTplus, Grant Agreement No. 754411. YIL acknowledges funding from the European Union’s Horizon 2020 research and innovation programme under the Marie Skłodowska-Curie Grant Agreement No. 101034413. The research was supported by funding to C-PH from the NOMIS Foundation, Project ID 1.844. The authors would like to thank past and present members of the Heisenberg and Hannezo groups for discussions, and, in particular, Shayan Shamipour, Viraj Doddihal, Michaela Jovic, Naoya Hino, Feyza Nur Arslan, Roksolana Kobylinska, and Carolina Camelo for feedback on the manuscript draft. This research was supported by the Scientific Service Units (SSU) of ISTA through resources provided by the Aquatics Facility, Imaging & Optics Facility (IOF), Scientific Computing (SciComp) facility, and Lab Support Facility (LSF).

## Author Contributions

NM and CPH conceptualized the study. All authors contributed to the design of the experiments. NM (experiments) and YIL (theory) acquired and analyzed the data. All authors contributed to manuscript preparation.

---

**Fig. S1:**
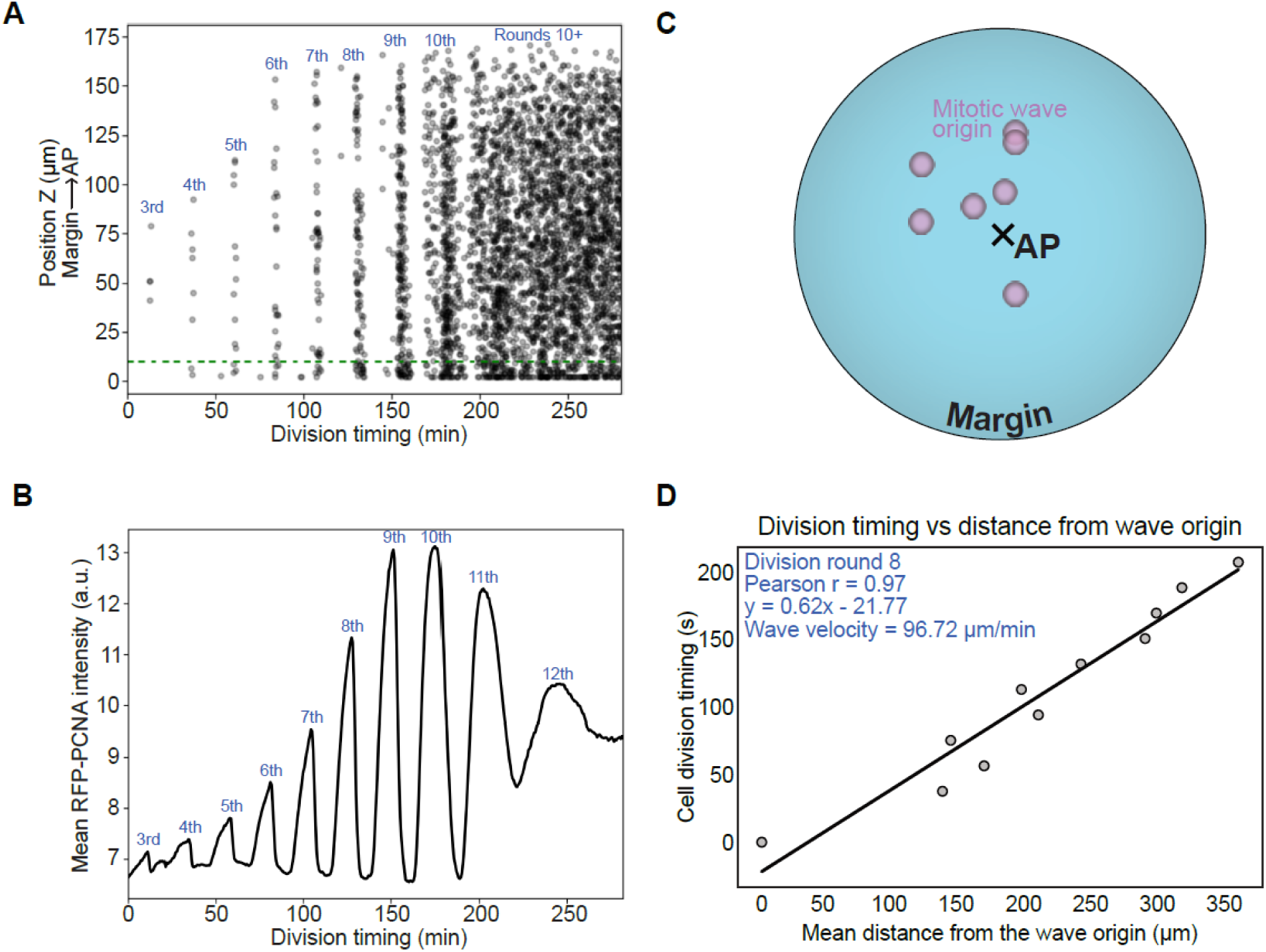
Cells cycle periodically, and their division timings correlate linearly with their distance from the wave origin. (A) Scatter plot of division timing for individual cells starting at cleavage division round 3 (x-axis) relative to their position along the animal-margin axis (y-axis) for one representative wild-type embryo. The green dotted line outlines cells located at a distance of <10 μm from the bottom of the image stack that could not be reliably tracked. (B) Mean RFP-PCNA intensity, as a marker for mitosis onset, as a function of time. Mean PCNA intensity increases during the S-phase, peaks just before the S-M transition, and declines in the M-phase. Thus, a narrow peak indicates greater synchrony and vice versa. (C) Representative scheme showing the position of the mitotic wave origin in division round 8 (n=7 embryos). (D) Representative plot of cleavage division timing as a function of cell distance from the mitotic wave origin in cleavage division round 8 from a wild-type embryo. Cells undergoing the S-M transition were binned together.

**Fig. S2:**
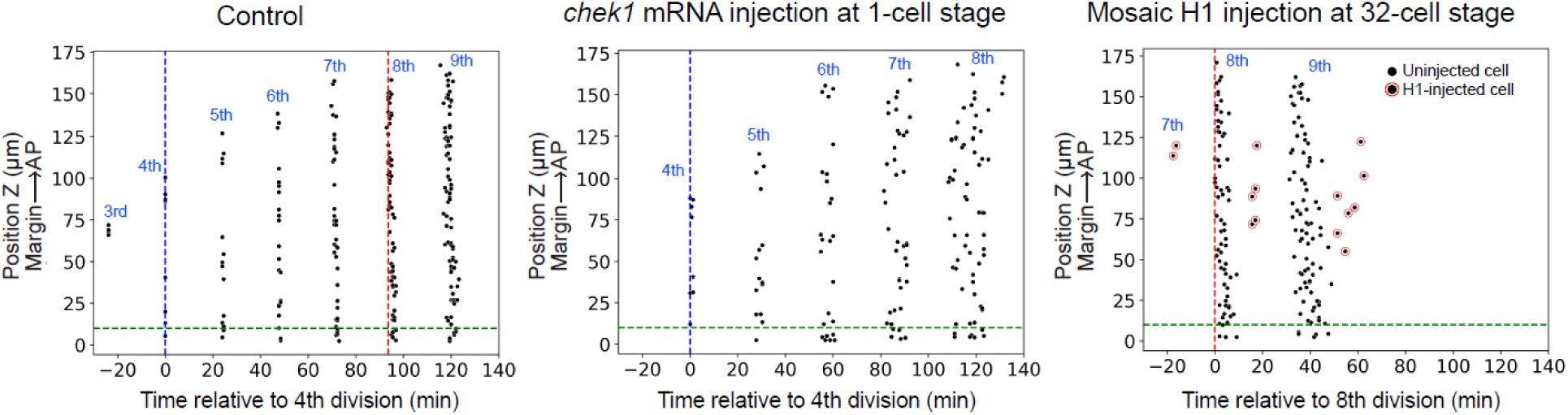
Desynchronized cell cycles do not resynchronize during cleavages. Representative scatter plots of the division timing for individual surface cells relative to their position along the animal-margin axis in control embryos (left), and embryos injected with either 12pg *chek1* mRNA at the 1-cell stage (middle) or 0.3pg Histone1 protein at the 32-cell stage (right). Blue and red dotted lines demarcate the onset of the 4th and 8^th^ cleavage divisions, respectively. Green dotted line outlines cells located at a distance of <10 μm from the bottom of the image stack that could not be reliably tracked.

**Fig. S3:**
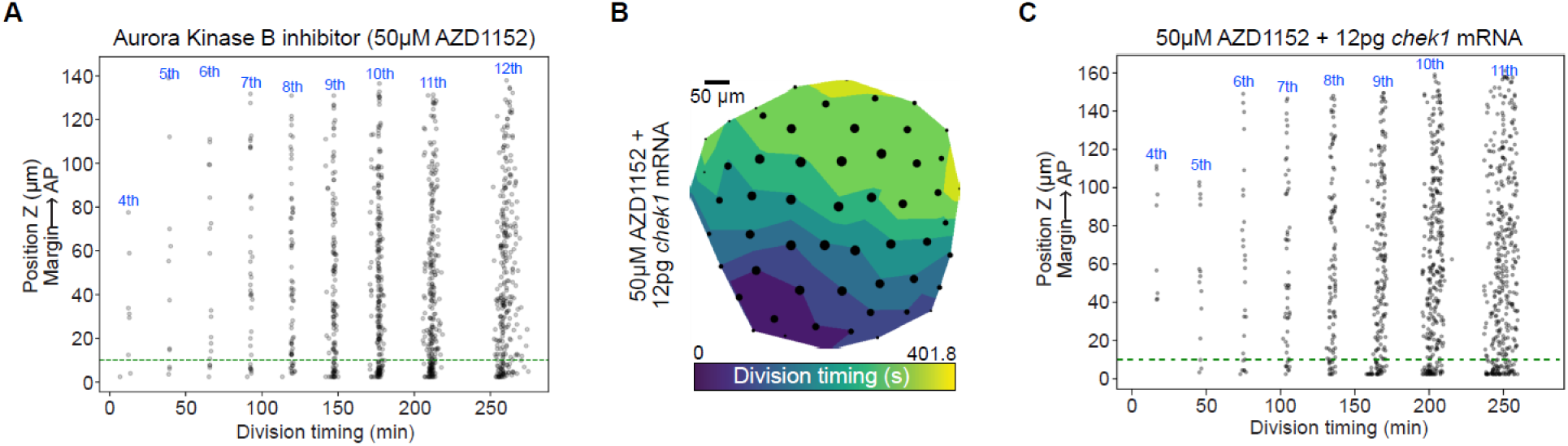
Mitotic wave originates at the margin and persists after division round 10 in syncytial embryos (A) Representative scatter plot of the division timings for individual cells relative to their position along the animal-margin axis in a syncytial Tg(*actb2:rfp-pcna*) embryo (n=7 embryos). (B) Contour plot of the mitotic wave across all surface cells at the 8^th^ round of cleavage division round in syncytialized Tg(*actb2:rfp-pcna*) embryos injected with 12pg *chek1* mRNA at the one-cell stage, and (C) a scatter plot of division timings relative to position along the AP-margin axis for a representative syncytial embryo injected with 12pg *chek1* mRNA (n=4 embryos).

**Fig. S4:**
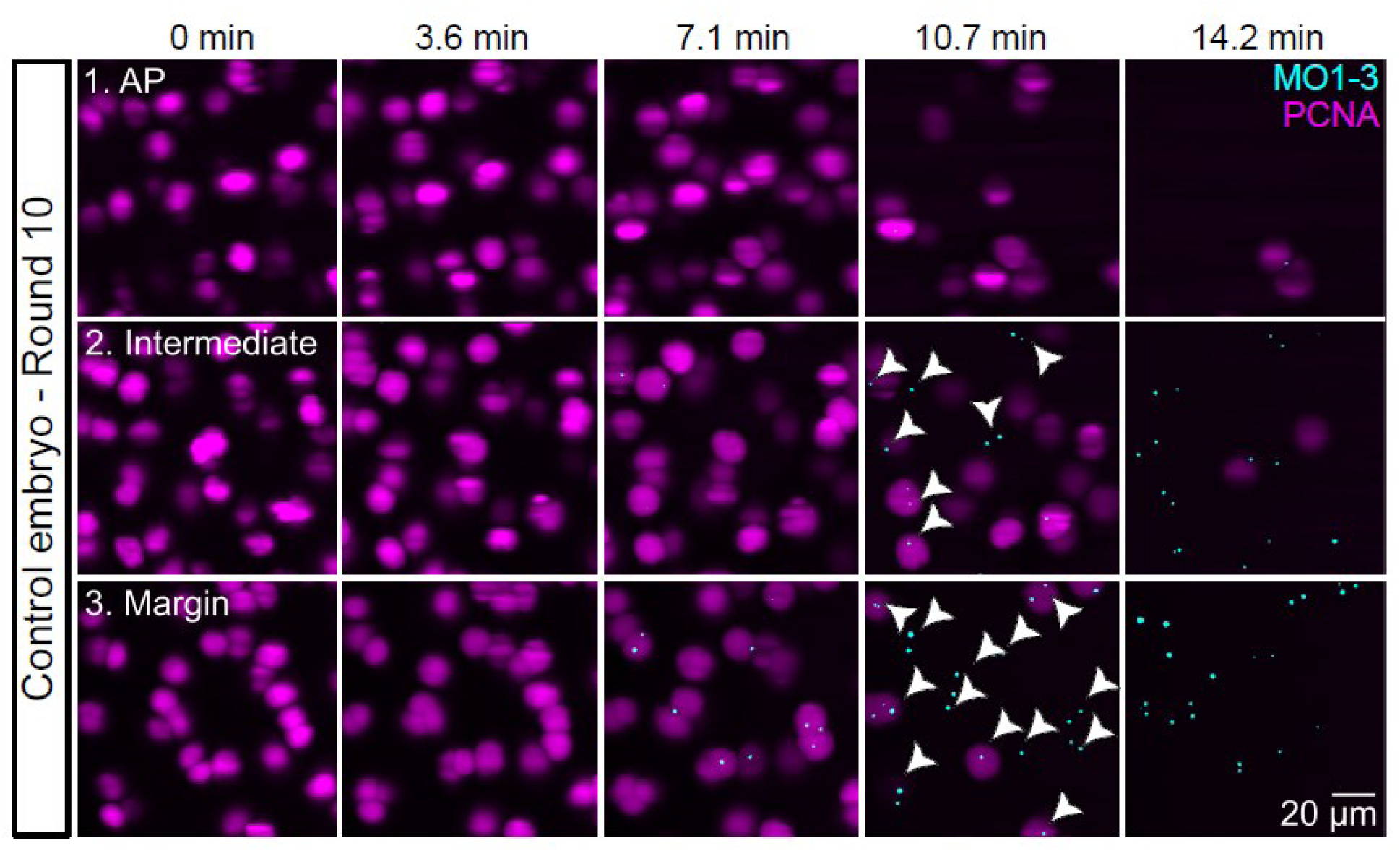
Zygotic transcription is initiated in a patterned manner along the AP-margin axis Zoomed-in views of the insets (1, animal pole; 2, intermediate region; 3, margin) shown in Fig. 5A for a control Tg(*actb2:rfp-pcna)* embryo injected with MO1-3-Fluorescein in the 10^th^ division round. Cyan, MO1-3-Fluorescein (nascent *mi430* transcripts); magenta, nuclei (PCNA). Each inset represents an area of 120 µm X 120 µm in size.

## Movie legends

**Movie S1:** Animal pole (AP)-view of the cell cycle progression beginning from division round 3 in a Tg(*actb2:rfp-pcna*) embryo. Temporal resolution: 38.7 seconds/frame.

**Movie S2:** AP-view of the cell cycle progression beginning from division round 7 in a Tg(*actb2:rfp-pcna*) embryo with a mosaic injection of 0.3pg Histone 1-Alexa Flour 488 at the 32-cell stage. The highlighted region indicates the position of the (daughters of the) injected cells. Red: PCNA, Green: Histone 1. Temporal resolution: 39.1 seconds/frame.

**Movie S3:** AP-view of the cell cycle progression beginning from division round 4 in a Tg(*actb2:rfp-pcna*) embryo injected with 12pg *chek1* mRNA at the one-cell stage. Temporal resolution: 37.9 seconds/frame.

**Movie S4:** AP-view of the nuclear cycling beginning from division round 4 in a Tg(*actb2:rfp-pcna*) embryo syncytialized by inhibiting Aurkb activity through 50µM AZD1152-treatment at the one-cell stage. Temporal resolution: 38.1 seconds/frame.

**Movie S5:** AP-view of the nuclear cycling beginning from division round 3 in a syncytialized Tg(*actb2:rfp-pcna*) embryo injected with 12pg *chek1* mRNA at the one-cell stage. Temporal resolution: 57.4 seconds/frame.

**Movie S6:** AP-view of the cell cycle progression beginning from division round 4 in a Tg(*actb2:rfp-pcna*) embryo from which one nucleus at the two-cell stage was aspirated. (Top: Control half, Bottom: Nucleus aspiration half). Temporal resolution: 71.8 seconds/frame.

**Movie S7:** AP-view of the cell cycle progression beginning from division round 4 in a bi-lobed Tg(*actb2:rfp-pcna*) embryo. Temporal resolution: 66.7 seconds/frame.

**Movie S8:** AP-view of the cell cycle progression beginning from division round 5 in a Tg(*actb2:rfp-pcna*) embryo with the yolk-blastoderm curvature increased through a partial severing of the yolk at the two-cell stage. Temporal resolution: 65.3 seconds/frame.

**Movie S9:** AP-view of the cell cycle progression beginning from division round 3 in a Tg(*actb2:rfp-pcna*) embryo treated with 10µM PF477736 to inhibit Chek1 activity. Temporal resolution: 65.9 seconds/frame.

**Movie S10:** AP-view of the cell cycle progression and *mirR430* pri-mRNA beginning from division round 8 in a control Tg(*actb2:rfp-pcna*) embryo injected with MO1-3-Fluorescein at the one-cell stage. Magenta: PCNA, Cyan: MO1-3. Temporal resolution: 53.4 seconds/frame.

**Movie S11:** Lateral view of the cell cycle progression and *mirR430* pri-mRNA beginning from division round 3 in a control Tg(*actb2:rfp-pcna*) embryo injected with MO1-3-Fluorescein at the one-cell stage. Red: PCNA, Green: MO1-3. Temporal resolution: 60 seconds/frame.

**Movie S12:** AP-view of the cell cycle progression and *mirR430* pri-mRNA beginning from division round 8 in a bi-lobed Tg(*actb2:rfp-pcna*) embryo injected with MO1-3-Fluorescein at the one-cell stage. Magenta: PCNA, Cyan: MO1-3. Temporal resolution: 66.7 seconds/frame.

