## Supplementary material for "Geometry-driven asymmetric cell divisions pattern cell cycles and zygotic genome activation in the zebrafish embryo": Detailed description of the theoretical modelling and analysis of cell cycle synchronization.

In this supplementary information, we demonstrate how spatial gradients, boundary conditions, and cell-cell cooperation interact to give rise to complex mitotic wave patterns, using the Kuramoto model of coupled oscillators. We derive a phase diagram and demonstrate how this can explain the diverging phenomenology observed in intact and syncytial embryos.

### T1. KURAMOTO MODEL ON A HEMISPHERE

The Kuramoto model [1] describes how interacting oscillators can spontaneously synchronize, and has been widely applied to various biological systems, from circadian clocks [2] to somitogenesis [3–5], to spermatogenesis [6]. In our context, each oscillator represents a cell of the zebrafish embryo, with its phase ($\theta$) representing the cell's state in the cell cycle. Cells that are physically adjacent can influence each other's timing through various signaling mechanisms, which we represent through local coupling between neighboring oscillators.

To capture the geometry of the zebrafish embryo and distribute cells on the hemisphere, a natural approach might be to use a regular latitude-longitude grid, but this would create artificial density variations near the poles. Similarly, projections of regular polyhedra onto the sphere introduce systematic biases at their vertices and edges. Instead, we arrange $N$ oscillators on a hemispherical Fibonacci lattice [7] (Figure T1), which provides several key advantages: (i) it provides the most uniform point distribution achievable by any deterministic algorithm; (ii) it maintains consistent nearest-neighbor distances across the surface, matching the approximately uniform cell sizes observed in experiments; (iii) it avoids special points or preferred directions that could introduce artificial biases in simulations.

The phase of each cell's cycle evolves according to:

$$\partial_t \theta_i = \omega_i + \epsilon \sum_{j \in \mathcal{N}_i} \sin(\theta_j - \theta_i) \tag{T1}$$

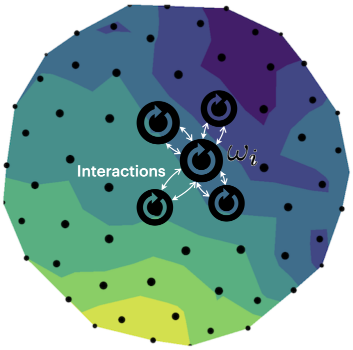


FIG. T1. Schematic representation of our model. Each black dot represents a cell positioned on a hemispherical Fibonacci lattice. Dot size increases with height ($z$) to aid visualization in this top-down projection. White arrows between dots indicate coupled neighbors that can influence each other's cell cycle timing.

Here, $\theta_i$ represents the cell cycle phase of cell $i$, and $\mathcal{N}_i$ denotes its nearest neighboring cells, determined through Delaunay triangulation of the hemisphere. The coupling strength $\epsilon$ represents how strongly each cell influences its neighbors' timing. The intrinsic frequency $\omega_i$ represents how quickly cell $i$ would progress through its cell cycle in isolation.

From wild-type measurements, we observed that cell cycle periods vary systematically with height ($z$) in the embryo according to:

$$T = T_0(1 + kz/L) \tag{T2}$$

where $T_0$ is the base period, $k = 1.9\%$ is the measured linear coefficient (percentage difference between the period of cells at the animal pole and the period of cells at the margin), and $L$ is the overall height of the hemisphere. We therefore set each oscillator's intrinsic frequency as $\omega_i = 2\pi/T_i(z) + \Lambda_i$, where $\Lambda_i$ represents biological noise. This noise term follows a normal distribution with mean 0 and standard deviation $\sigma$, with different cells' noise terms being independent: $\langle \Lambda_i \Lambda_j \rangle = \delta_{ij}\sigma^2$.

### T2. STEADY-STATE PHASE DIAGRAM

Clearly, in the absence of any coupling or noise, waves are deterministic and simply follow the gradient in periods imposed in the system (i.e. waves arising from the anterior pole, being slow and slow as a function of time, following the theoretical expression discussed in the main text and observed experimentally in Fig. 1). However, in the presence of either noise or coupling, more complex behaviors could be observed. In analyzing the behaviors of the model, we find that the final pattern of cell division times depends on the balance between two competing factors: how strongly cells communicate with their neighbors (represented by $\epsilon$) and how much random variation exists in their natural division times (represented by $\sigma$). As shown in Fig T2a, with small noise, the wave pattern follows the pre-determined AP-centric pattern of the periods; as the noise increases, if the interaction strength is sufficient to compensate for the increase in noise, margin-originated waves are observed, otherwise, the phases are desynchronized.

Furthermore, we can show that if the waves do emerge spontaneously, the speed of the wave is also solely determined by $r = \epsilon/\sigma$. We can demonstrate this mathematically by rescaling time in our original equation. If we introduce a new time variable $t' = \epsilon t$ (which gives $\partial_t = \epsilon\partial_{t'}$), after relabelling $t'$ back to $t$, our equation becomes:

$$\partial_t \theta_i = \frac{1}{\epsilon}\omega_i + \sum_{j \in \mathcal{N}_i} \sin(\theta_j - \theta_i) \tag{T3}$$

This rescaling reveals that the system's long-term behavior depends only on the ratio $r = \sigma/\epsilon$. When this ratio is small (strong coupling relative to noise), cells tend to coordinate their divisions into clear fast waves. As the ratio increases, the waves tend to be slower. When the ratio is very large (weak coupling relative to noise), the system becomes disordered, with cells dividing independently. This behavior is shown in the phase diagram in Figure T2b: lines of constant ratio $r$ approximately correspond to the same variance in division times in the steady state.

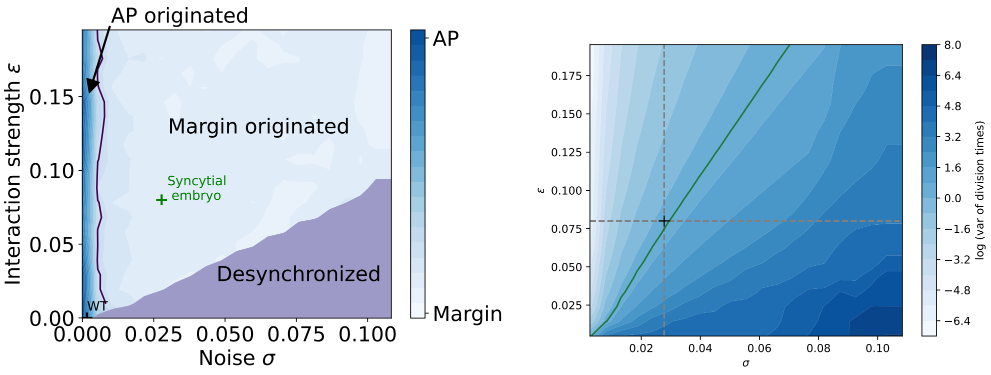


(a) origin of waves

(b) phase diagram

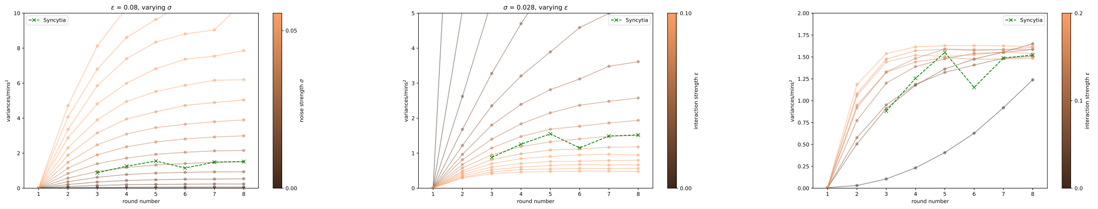


(c) fixed $\epsilon$

(d) fixed $\sigma$

(e) variances over time

FIG. T2. (a) Origin of the waves in $\epsilon - \sigma$ space. The boundary between AP originated and margin originated is thresholded at $z = 0.5$ and the region of desynchronization corresponds to $\langle e^{i\theta_j} \rangle_j < 0.6$, where $\langle \rangle_j$ denotes averaging over all sites [1] (b) Phase diagram of the Kuramoto model in $\epsilon - \sigma$ space. The log of the variance of the division times at the 8th round is plotted. The green solid line indicates the contour line of the observed variance at the 8th round in syncytia embryos. The black cross indicates the best-fit parameter for the syncytia embryo. (b) Variances of division times for each round for fixed $\epsilon$ as $\sigma$ varies. Parameters are taken along the horizontal dashed line in the phase diagram. (c) Variances of division times for each round for fixed $\sigma$ as $\epsilon$ varies. Parameters are taken along the vertical dashed line in the phase diagram. (d) Variances of division times over time for fixed $\epsilon/\sigma$ ratio (along the green contour line in the phase diagram) as both parameters are varied proportionally. Although the final variance is the same for simulations, the time it takes to reach this variance decreases with increasing $\epsilon$.

### T3. TIME EVOLUTION DYNAMICS

While the ratio between cell-cell communication strength ($\epsilon$) and biological noise ($\sigma$) determines the final pattern of division waves, we need to understand how quickly these patterns emerge to fully match our experimental observations. This temporal evolution helps us determine the actual values of $\epsilon$ and $\sigma$, not just their ratio.

We tracked how variable the division times were across the embryo for the first eight cell division cycles (Figure T2e). As discussed above, in the simplest case, with no cell-cell communication ($\epsilon = 0$), these timing differences, regardless of source, accumulate with each round of division, causing the variance to increase quadratically over time. When we introduce cell-cell communication ($\epsilon > 0$), neighboring cells begin to influence each other's timing. If this communication is strong enough, it can overcome the natural gradient in cell cycle periods, leading to more synchronized divisions. The stronger the communication (larger $\epsilon$), the faster this synchronization occurs.

Importantly, this phenomenology could describe well the syncytium data, showing a plateau in variance after a

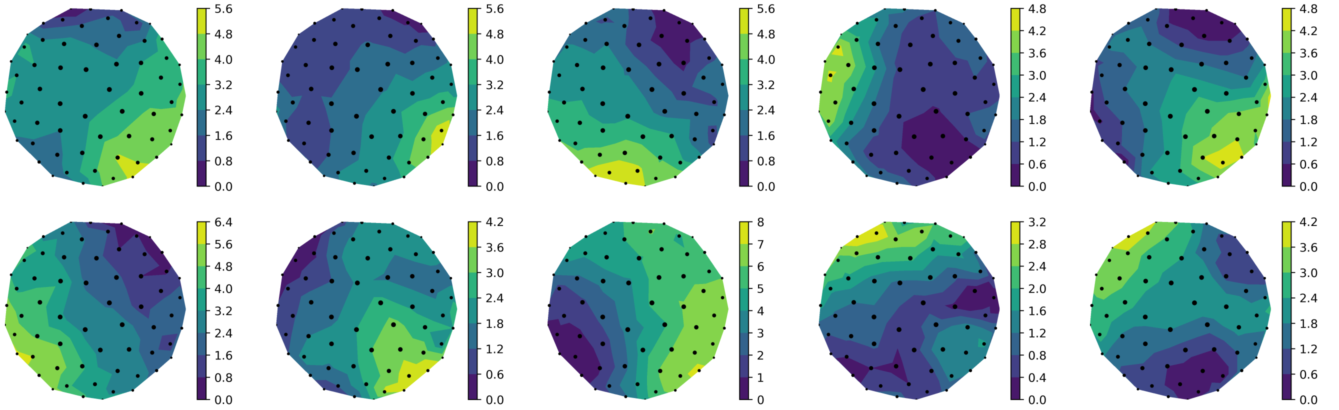


FIG. T3. 10 realizations of the model with $\epsilon = 0.08, \sigma = 0.028$. The division times of the 8th round are shown. Despite some variation between simulations, all realizations show similar wave-like patterns of division times originating at the margin.

few cleavages. By comparing various points along the black contour line in Figure T2b to our experimental data from syncytial embryos, we found that the model best matches the data when $\epsilon = 0.08$ and $\sigma = 0.028$. These values capture both the final pattern of division times and how quickly the pattern emerges. To demonstrate the robustness of our model, Figure T3 shows ten different simulations using these parameters, focusing on the division times at the eighth round of cell division.

### T4. THEORY FOR WT EMBRYOS

Having established a framework for understanding division waves in syncytial embryos, we now extend our analysis to intact WT embryos. As discussed in the main text, experimental perturbations point to limited couplings in intact embryos. From a theoretical perspective, in the absence of any coupling ($\epsilon = 0$), the variance in division times would increase quadratically with each cell cycle due to the accumulation of timing differences, with the slope of this increase dependent on noise levels. However, our experimental measurements of variance in WT embryos (Figure T4) reveal a more complex pattern: the variance increases more slowly in early rounds compared to later rounds. One possibility to explain this would be that the intrinsic noise level $\sigma$ increases as a function of developmental time. Another is that at early stages, cell-cell communication might be still present, a noise-buffering mechanism that decays over developmental time.

As discussed in the main text, given previous reports that cytokinesis in the early embryo is incomplete, leading to intercellular bridges that resolve over developmental time, the most parsimonious hypothesis is simply that cell-cell couplings are proportional to bridge size – and thus that at very early stages, WT model parameters are closer to syncytium, while at later stages, coupling is essentially zero. Assuming that the size of the intercellular bridges decreases exponentially with time as cells divide, the following function well captures the interaction strength: $\epsilon(t) = \epsilon_0 e^{-n}$, where $\epsilon_0$ is the initial coupling strength and $n$ is the number of rounds since the first division. Setting $\epsilon_0 = 0.08$ (matching the coupling strength found in syncytial embryos) and fitting $\sigma$ to the experimental data yields good agreement with the observed pattern of variances (Figure T4) up to the 8th round of division.

Interestingly, beyond the 9th round, as shown in Figure T4b, the variance exhibits an upward trend that is much

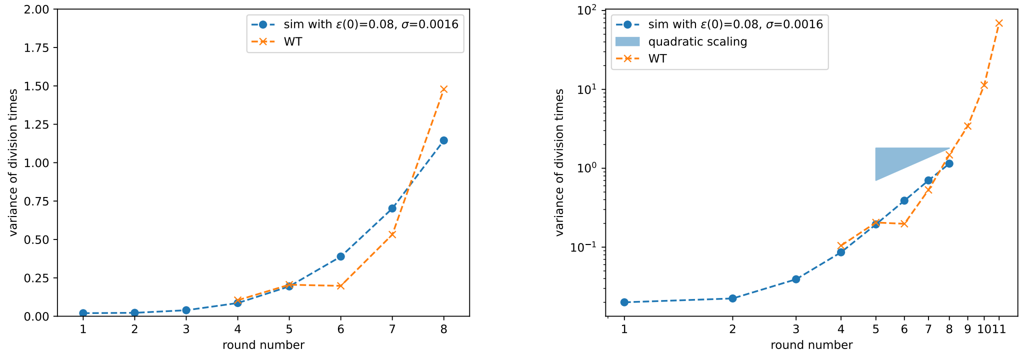


(a) WT fit on linear scale (b) WT fit on log-log scale

FIG. T4. Variances of division times for each round in WT embryos, on linear and log-log scales.

steeper than quadratic scaling, something that even the non-interacting scenario cannot capture. This means that the noise in the system must also increase drastically in the later rounds, i.e. at the onset of midblastula transition, which requires further investigation and is beyond the scope of the current paper.

---

[1] Y. Kuramoto. *Chemical Oscillations, Waves, and Turbulence*. Springer, 1984.
